# Timing and Amplitude of Catch-up Saccades to Accelerating Targets

**DOI:** 10.1101/2024.01.11.575233

**Authors:** Sydney Doré, Jonathan Coutinho, Emma Brooks, Aarlenne Z Khan, Philippe Lefèvre, Gunnar Blohm

**Author notes:** Corresponding Author: Dr. Gunnar Blohm, Centre for Neuroscience Studies, Queen’s University, 18, Stuart Street, Kingston, ON, K7L 3N6, Canada.

## Abstract

To track moving targets, humans move their eyes using both saccades and smooth pursuit. If pursuit eye movements fail to accurately track the moving target, catch-up saccades are initiated to rectify the tracking error. It is well known that retinal position and velocity errors determine saccade timing and amplitude, but the extent to which retinal acceleration error influences these aspects is not well quantified. To test this, 13 adult human participants performed an experiment where they pursued accelerating / decelerating targets. During ongoing pursuit, we introduced a randomly sized target step to evoke a catch-up saccade and analyzed its timing and amplitude. We observed that retinal acceleration error was a statistically significant predictor of saccade amplitude and timing. A multiple linear regression supported our hypothesis that retinal acceleration errors influence saccade amplitude in addition to the influence of retinal position and velocity errors. We also found that saccade latencies were shorter when retinal acceleration error increased the tracking error and vice versa. In summary, our findings support a model in which retinal acceleration error is used to compute a predicted position error ∼100ms into the future to trigger saccades and determine saccade amplitude.

**Significance statement:** When visually tracking object motion, humans combine smooth pursuit and saccadic eye movements to maintain the target image on the fovea. Retinal position and velocity errors are known to determine catch-up saccade amplitude and timing, however it is unknown if retinal acceleration error is also used to predict future target position. This study provides evidence of a small but statistically significant contribution of retinal acceleration error in determining saccade amplitude and timing.

## Introduction

Humans utilise two complementary eye movement types to track moving objects - smooth pursuit and saccades. Smooth pursuit is primarily driven by visual motion, but occasionally, tracking errors can accumulate, necessitating a catch-up saccade to realign the gaze with the target and maintain foveation. The mismatch between the eye and target during tracking results in a position error that is used to determine the timing and amplitude of the resulting catch up saccade (Rashbass, 1961; Robinson 1986; Keller & Johnsen 1990; De Brouwer et al. 2002). Recently, Coutinho et al. (2021), and Nachmani et al. (2020) proposed that the eye movement system predicts position error ∼100ms into the future through motion extrapolation using position and velocity tracking errors. This predicted position error is used both to determine saccade timing and compute saccade amplitude. They proposed a predictive, probabilistic process that relies on the accumulation of noisy sensory position and velocity error signals towards a decision threshold to trigger a saccade. This noisy evidence is also used to determine the amplitude of the upcoming saccade.

It is known that in addition to velocity and position inputs, target acceleration is also used in the smooth pursuit system; however it is not well understood how acceleration affects saccades. In studies involving the tracking of sinusoidal target movements by both humans and monkeys, it was observed that eye acceleration correlated with velocity error, while smooth pursuit gain was influenced by target acceleration. Notably, higher maximum target accelerations were associated with lower pursuit gain (Lisberger, Evinger, Johanson & Fuchs, 1981). Results of this experiment suggest that information about target acceleration in addition to target speed is provided to the smooth pursuit system. Image motion models of smooth pursuit also use visual input signals related to target acceleration (Morris & Lisberger, 1985; Lisberger et al., 1987, Krauzlis & Lisberger, 1989, Krauzlis & Lisberger, 1994). In studies using transient occlusion, target acceleration influenced the control of pursuit before and during the occlusion, but the accelerating target needed to be presented for at least 500ms prior to the occlusion; eye velocity at the end of the occlusion was scaled to target acceleration (Bennett & Barnes 2006; Bennett, Orban de Xivry, Barnes & Lefevre, 2007; Bennett & Benguigui, 2013). Findings from these studies indicate that target acceleration can be extracted and used by the pursuit system, and this capability enhances with longer presentation times. More recently, other studies using occlusion tasks have shown that, while humans can accurately pursue accelerating targets as predictive pursuit scales with target acceleration, manual target interception is not influenced by target acceleration but instead uses an extrapolation of pre-occlusion velocity (Kreyenmeier, Kämmer, Fooken & Spering, 2021; 2022). Studies that have investigated the behavior of eye movements to perturbations in target velocity have also provided evidence for the use of target acceleration in influencing pursuit (Tavassoli and Ringach, 2010; Brostek et al., 2017). In sum, target acceleration is used to modulate pursuit responses. Given that the saccadic and pursuit systems are synergistic and share signals both at the neurophysiological (Krauzlis, 2004) and the behavioral level (Orban de Xivry & Lefevre, 2007), it follows logically that target acceleration could also be used by the saccadic system for catch-up saccade planning.

While previous studies have investigated the influence of occlusion, sinusoidal and circular target acceleration tracking with respect to saccades, an understanding of how constant target acceleration is used in computing catch-up saccades is lacking. In the occlusion study discussed above, saccadic eye displacement was modulated by target acceleration only when presented for at least 500 ms (Bennett et al., 2007). These researchers also found that saccadic eye displacement during transient occlusion changed proportionately to target acceleration under these conditions. The results of this study suggest that target acceleration information can be extracted and used to predict the occluded target’s trajectory. Another target occlusion study also investigated the influence of accelerating target motion on predictive saccades (Kreyenmeier, Kämmer, Fooken & Spering, 2022). Different levels of target acceleration were not shown to have scaled with the landing time of the predictive saccade after temporal occlusion, however saccades landed later for accelerating motion than they did for decelerating motion (Kreyenmeier, Kämmer, Fooken & Spering, 2021). The landing times of these predictive saccades suggest that target acceleration was not taken into account during occlusion, and instead were computed based on the last available velocity information pre-occlusion. Although participants’ predictive saccades landing times did not scale to target acceleration, it is unknown how target acceleration affected saccade amplitude or timing in this study. Furthermore, as these were predictive saccades during an occlusion period, they do not reflect the behavior of saccades when tracking uninterrupted target motion.

The purpose of the current study is to investigate whether participants use retinal acceleration error in computing the timing and amplitude of catch-up saccades during ongoing smooth pursuit. We hypothesized that retinal acceleration error would be a significant predictor for both the timing of catch-up saccades and the computation of catch-up saccade amplitudes. We expect that retinal acceleration error will modulate catch-up saccade latencies depending on how retinal acceleration changes the predicted position error. We also predict that acceleration error will be a significant regressor along with velocity and position error in determining saccade amplitude. We tested these hypotheses in the context of targets that are continuously changing speed to ensure enough time for the brain to estimate target acceleration. We then introduced a sudden position step (jump) in the motion trajectory of the target to trigger a catch-up saccade.

## Material & Methods

### Sampling Plan & Participants

The study’s procedures were approved by the Queen’s University General Research Ethics Board in accordance with the Declaration of Helsinki. A power analysis based on effect sizes from previous work done in our lab (Nachmani et al., 2020) indicated the minimum number of participants needed is 12. To anticipate attrition, 15 adult participants were recruited from the Queen’s University Centre for Neuroscience Studies graduate pool. Participants provided informed consent and received compensation at a rate of $10 per hour for their involvement. To qualify for participation, individuals needed to be at least 18 years old and possess either normal vision or corrected-to-normal vision. Two participants opted to withdraw from the study, resulting in a final analysis sample size of 13 participants. This group had an average age of 21 years, comprising 8 females and 5 males.

### Experimental Procedure

We used a previously established double step-ramp task (Rashbass, 1961; de Brouwer et al., 2002) generated using custom Matlab code (Mathworks, Inc), with the Psychophysics Toolbox (Brainard, 1997) into which we introduced a new acceleration component. This task involved an abrupt change in target position used to trigger catch–up saccades. Stimuli were displayed on a ViewPixx screen (VPixx Technologies, 120Hz refresh rate, resolution 1920 x 1200, strobed backlight) positioned 50 cm away from the participant spanning 60 degrees of their visual field. Eye movements from the right eye were recorded using an Eyelink 1000 video-based recording system (SR Research, Mississauga, ON, Canada) at 1000 Hz. Participants underwent a standard 9 dot calibration (Eyelink) every 3 blocks to ensure accurate eye position recording. Trials were configured in one of two ways, either an accelerating or decelerating trial, with an initial fixation target positioned 20 degrees to either the left or right side of the visual field. The visual stimulus was a white dot on a black screen that moved horizontally. Target acceleration was a random integer variable between −80 to 80 deg/s^2^ (excluding 0). Regardless of the initial fixation target position, two scenarios would occur as described in Figure 1. Accelerating trials began with an initial velocity of 0, continuously accelerating in the direction opposite to the fixation position after a 750 –1250 ms fixation period. After a random period between 300-500 ms, the target jumped (position step) randomly between −6 and 6 degrees, while target acceleration remained the same for another 500-700 ms, followed by a 500 ms fixation period. Decelerating trials consisted of a first position step of 6 degrees and a starting velocity of −40 deg/s if the initial fixation was positive (right side of screen), and −6 degrees and a velocity of 40 deg/s if the initial fixation was negative (left side of screen). Thereafter, an additional position step randomly chosen between −6 and 6 degrees occurred after 300-500 ms with no change in target acceleration, followed by another 500-700 ms and ending with a 500 ms fixation period. These initial fixation positions, initial target velocities, position steps and acceleration values varied randomly between each trial. (Note: rightward positions are positive values)

**Figure 1:**
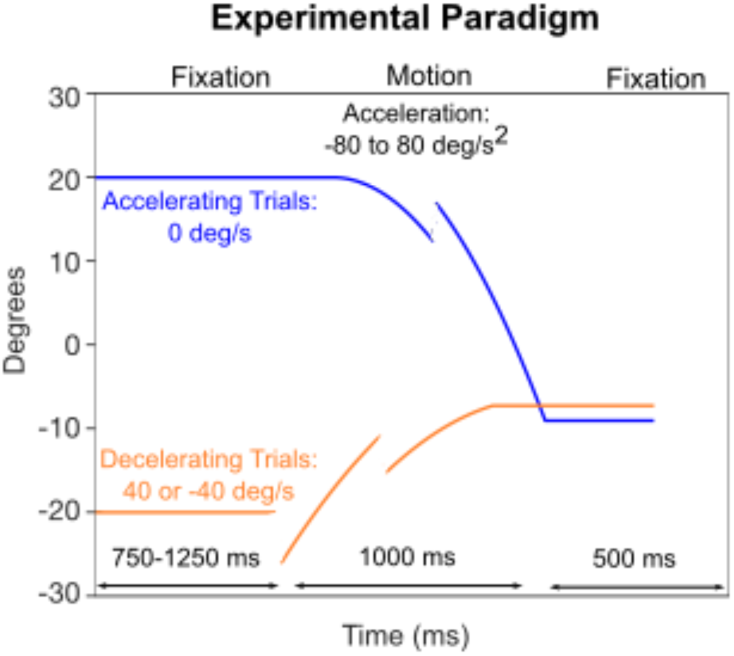
Experimental paradigm. The blue trace depicts an example trial with an accelerating target; the orange trace shows an example trial with a decelerating target trajectory. Trials started with a 20 deg left or right initial fixation target. Target acceleration was a random integer variable between −80 to 80 deg/s^2^ (excluding 0). Accelerating trials (blue) began with an initial velocity of 0 deg/s, and continuously accelerated in the direction opposite to the fixation position after a 750 –1250 ms fixation period. Decelerating trials consisted in a 6 deg outward target step (away from the screen center) and a starting velocity of 40 deg/s toward the center of the screen. For both trial types, after a random period between 300-500 ms, the target jumped (position step) randomly between −6 and 6 deg, while target acceleration/deceleration remained the same for another 500-700 ms, followed by a 500 ms fixation period.

### Data Pre-processing

Each participant undertook 5 data collection sessions of approximately 30 minutes each. Each session consisted of 10 blocks each with 50 trials, resulting in 2500 trials per participant and an overall total of 32,500 trials. Eye position was low-pass filtered using an auto-regressive forward-backward filter with a cut-off frequency of 50Hz. Eye velocity and acceleration were derived from position signals using a central difference algorithm (±10ms window), and saccades were detected using an acceleration threshold of 750 deg/s^2^. We visually inspected all trials for errors using a custom-made analysis interface in Matlab. Trials in which the target was not tracked by the participant (e.g. due to distraction), that contained blinks between 100 ms before the step and the first catch-up saccade, that consisted of catch-up saccades where there was more than one velocity peak, saccades which occurred during the target jump during target motion, or trials that were missing eye tracking data were discarded from the analysis. Once data was extracted and analysed using Matlab, trials that had catch-up saccades with a latency relative to the target jump of less than 90 ms were also discarded from the analysis. 7678 trials were discarded from the analysis, with 24,822 remaining (76.4% of total trials). Each participant had between 1089 and 2269 trials with a mean of 1904 trials each.

### Eye Movement Parameters

The saccades of interest were the first saccades that occurred after the target step during target motion. It is well-established that visual stimuli do not exert any influence on saccades during the ∼100 ms period immediately preceding saccade onset (Orban de Xivry & Lefèvre, 2007). Therefore, all statistical analyses pertaining to saccade amplitude were conducted using data from the 100 ms timepoint prior to the onset of the saccade. Position errors were determined by subtracting the eye’s position 100 ms before the saccade initiation from the corresponding target position at that moment. Velocity errors (i.e., retinal slip) were computed by subtracting eye velocity from target velocity averaged over a 50ms window centered on 100 ms before the saccade. Acceleration error was computed using the slope of a robust linear regression of the eye velocity with the saccadic component removed computed over a a 200ms range before the saccade (centered at 100ms before the saccade). Both velocity and acceleration error computations used averaging over windows that included information less than 100 ms before the saccade in order to obtain an accurate estimate of the signals of interest from noisy data. Saccade amplitude was computed by subtracting the eye position at the beginning of the saccade from the eye position at the end of the saccade. Following the methods described by de Brouwer et al. (2002), saccade amplitude was then corrected to remove the smooth pursuit component; saccade duration and pursuit velocity during the saccade were multiplied and subtracted from the total amplitude to correct for the contribution of pursuit.

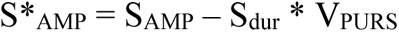

For all statistical analyses related to saccade timing, we evaluated the same variables at the time of the target step rather than 100 ms before the saccade. This was because saccade timing is always relative to the time of the step and thus sensory variables at the step predominantly determine saccade timing (de Brouwer et al., 2002; Nachmani et al., 2020; Coutinho et al., 2021).

As has previously been suggested, a predicted position error extrapolated to some time interval into the future is correlated with saccade timing and amplitude (Nachmani et al., 2020; Coutinho et al., 2021). Instantaneous predicted position error was thus calculated as:

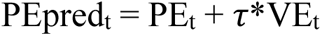

where retinal position error (“PE”) is the difference between target and eye positions and retinal velocity error (“VE”) is the difference between target and eye velocities. The variable “t” represents sampling time and “*τ*” is the extrapolation duration.

### Hypotheses

For our first hypothesis, we expected retinal acceleration error to be a significant factor involved in computing catch-up saccade amplitude. If retinal acceleration error was taken into account by the saccadic system, saccade amplitude should increase or decrease accordingly to ensure more accurate tracking. Thus, we assessed the influence of retinal position, velocity, and acceleration errors in computing catch-up saccade amplitude using a multiple linear regression analysis, similar to De Brouwer et al. (2002).

Our second hypothesis was that acceleration should also modulate saccade timing; retinal acceleration error should influence the error accumulation that is used to determine whether a saccade is triggered (Coutinho et al., 2021). Specifically, depending on the relative sign, retinal acceleration error would either increase or decrease the predicted position error, and thus saccade latencies should be shorter when the predicted position error is increased by retinal acceleration error and vice versa. This is because the same acceleration magnitude adding vs subtracting from the predicted position error should increase vs decrease the certainty with which a saccade is needed, thus leading to shorter vs longer saccade times, respectively. Therefore, we assessed the influence of estimated predicted position error on saccade timing. We used a repeated-measures ANOVA with the signed retinal acceleration error and predicted position error binned at four sizes as the independent variables and saccade latencies as the dependent variable.

As per our hypotheses, if retinal acceleration error was in fact used to time saccades and compute saccade amplitudes, then the equation for calculating predicted position error should be updated to include the additional acceleration error term as follows:

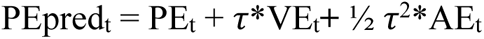

This effect of retinal acceleration error is expected to be small, even with large acceleration values, based on its contribution in the above equation.

## Results

Our study aimed to build on our current understanding of how saccade amplitude and timing is computed by assessing the influence of accelerating motion. Participants tracked a horizontal moving dot in an accelerating step ramp paradigm. Trials consisted of either accelerating (Figure 2A) or decelerating target motion (Figure 2D). As can be seen, in both conditions, participants were able to make fairly accurate catch-up saccades to foveate the target after the second target step. In panel A in the accelerating trial condition, corrected saccade amplitude (pursuit component removed; see Methods) was 11.75 deg while the position error 100 ms before the saccade was 9.76 deg (denoted by vertical double arrow). Saccade amplitude was similarly larger in magnitude than position error in the decelerating condition in panel D, with a corrected saccade amplitude of −5.27 deg and position error 100 ms before the saccade being −4.73 deg. Considering that the magnitude of position error was less than saccade amplitude, we can assume that velocity and/or acceleration errors 100 ms before the saccade were also taken into account when computing saccade amplitude (Hyp. 1), as we will later describe. Specifically, the accelerating trial condition displayed in Figure 2 had velocity and acceleration errors of 18.50 deg/s and 33.30 deg/s^2^ when sampled 100ms before the saccade, respectively. These same errors in the decelerating trial condition were −6.88 deg/s and 1.54 deg/s^2^. If these velocity and acceleration errors were not in fact taken into account, saccade amplitude should be the same size as the position error.

**Figure 2:**
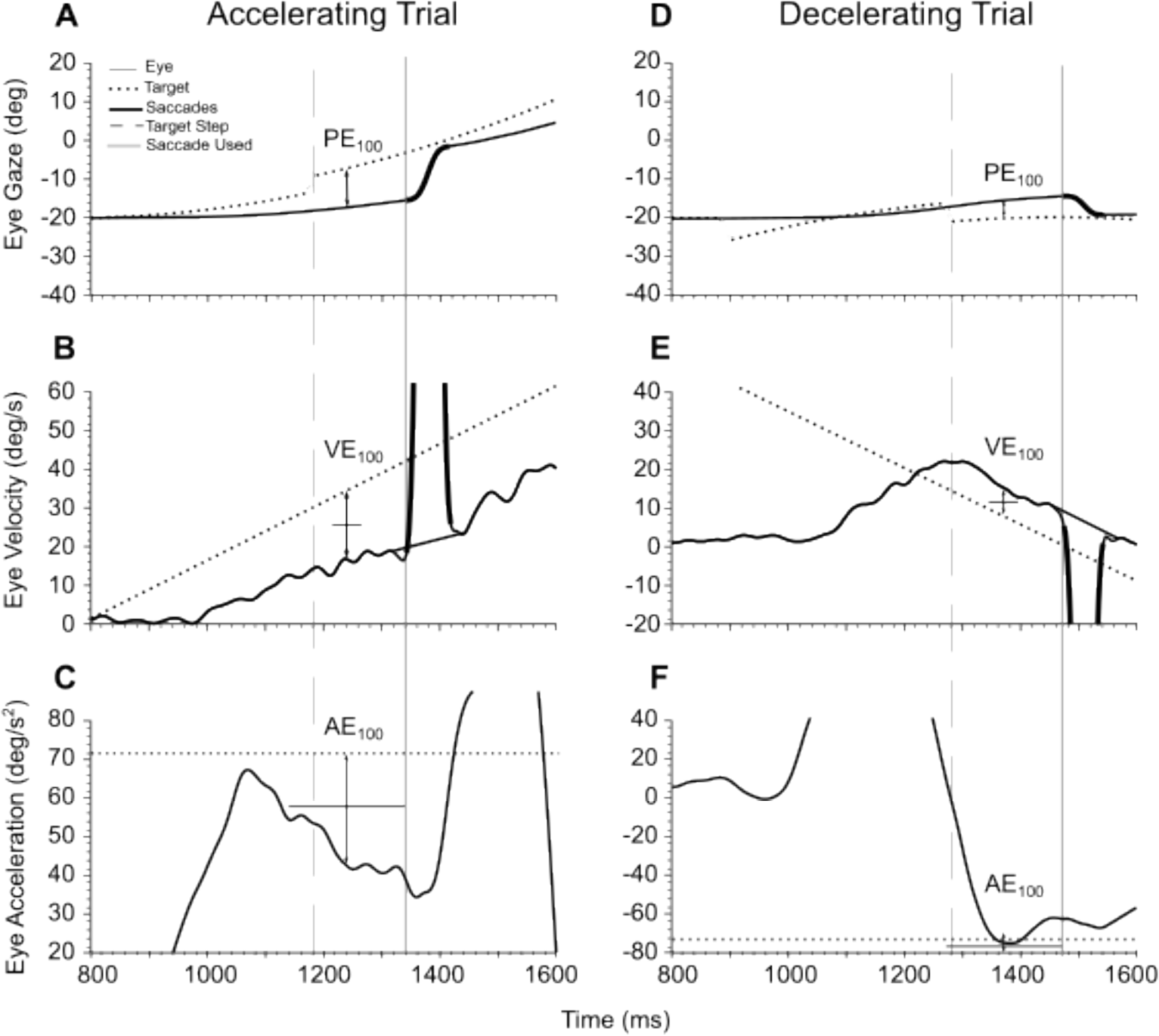
Typical trials. Example of a typical accelerating (left) and decelerating (right) trial. Eye and target position, velocity and acceleration respectively are plotted against time. Filled lines denote the eye, dashed lines denote the target. Bolded sections indicate a saccade. (A,D) Eye and target position, (B,E) velocity, and (C,F) acceleration vs time for (A, B, C) accelerating and (D, E, F) decelerating trial conditions.

Saccade latency was calculated as the time at which the saccade occurred relative to the target step, indicated by the grey vertical solid line (Figure 2). In the accelerating trial condition example (Fig. 2A), saccade latency was 160 ms. In the decelerating trial condition example (Fig. 2D), saccade latency was 190 ms. Predicted position error was computed using position and velocity error parameters at the target step (vertical grey dashed line), and is thought to be used to determine the timing of when the saccade is triggered relative to the target step (latency) which will be later described (Hyp. 2; see Catch-Up Saccade Timing section). Our aim was to determine if an acceleration component should be included in the computation of predicted position error. In Figure 2, position, velocity and acceleration errors at the target step in the accelerating trial condition example were 8.34 deg, 16.65 deg/s, and 31.09 deg/s^2^. Decelerating - 3.59 deg, −7.14 deg/s, and −71.47 deg/s^2^.

### Catch-up Saccade Amplitude

First, we wanted to examine whether retinal acceleration error was used to compute the amplitude of catch-up saccades. It is well known that retinal position and velocity errors ∼100 ms before saccade occurrence are used to compute saccade amplitude. We used a multiple linear regression to test if saccade amplitude also correlates with retinal acceleration in conjunction with position error and retinal slip sampled 100 ms before saccade onset. Saccade amplitude was corrected to remove the pursuit component (see Methods; Eye movement parameters section). All participants were included in the regression.

The fitted regression model was:

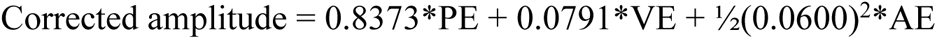

*Equation 1*.

The coefficients in eq 1. can be interpreted as follows. The coefficient of PE (0.8373) is the proportion of position error that is corrected by the saccade. The coefficient of VE (0.07910) is the time extrapolation duration that is multiplied by a given VE. The coefficient of AE (0.0018) can also be solved for the time extrapolation duration (0.0600), computed by solving for τ: β_AE_ =½(τ)^2^.

Retinal position error (PE), velocity error (VE) and acceleration error (AE) significantly predicted saccade amplitude (β_PE_ = 0.8373, p < 1.10^-100^; β_VE_ = 0.0791, p = p < 1.10^-100^; β_AE_ = 0.0018, p = 2.029724e-09; R^2^ = 0.893). See table 1 for more detail on regression results, including the standard errors and the upper and lower bounds on coefficients using a 95% confidence interval. The regression coefficients for VE and AE can be interpreted in terms of time extrapolation duration used to predict future target position, where VE = 79.1 ms and AE = 60.0 ms (see eq 1). With respect to the range of values we used for target position (−6,6 deg), velocity (−40, 40 deg/s) and acceleration (−80,80 deg/s^2^), we expected the relative contribution of acceleration to be small. For example, when computing saccade amplitude based on our coefficients in eq 1. using the maximum values in our parameter ranges, a 6 degree position error would result in a 5.024 degree correction, a 40 degree/s velocity error would result in a 3.164 degree correction, and a 80 degree/s^2^ acceleration error would result in a 0.144 degree correction.

**Table 1.** Multiple Linear Model Regression Results.

| Variable | Coefficient | Standard error | p value | 95% Confidence Interval |  |
| --- | --- | --- | --- | --- | --- |
|  |  |  |  | 0.025 | 0.975 |
| AE | 1.8e-03 | 0.3e-04 | 2.029724e-09 | 0.001 | 0.002 |
| VE | 7.9e-02 | 0.1e-03 | 0.000000e+00 | 0.077 | 0.081 |
| PE | 8.4e-01 | 0.2e-03 | 0.000000e+00 | 0.833 | 0.842 |

The scatterplot in figure 3 demonstrates predicted vs. observed saccade amplitude as each variable of interest is added for a single participant. As seen in figure 3a, the final addition of retinal acceleration error as a regression term tightens the relationship between predicted and observed amplitude. Figure 3b displays the regression residuals as each variable is added. Like figure 3a, it is evident from the visual representation that incorporating retinal acceleration error leads to the least amount of deviation from the diagonal line, i.e. the smallest difference between predicted and observed values. Standard deviations decrease and kernel density estimates tighten with the addition of each new variable, and is lowest when retinal acceleration error is included.

**Fig 3.**
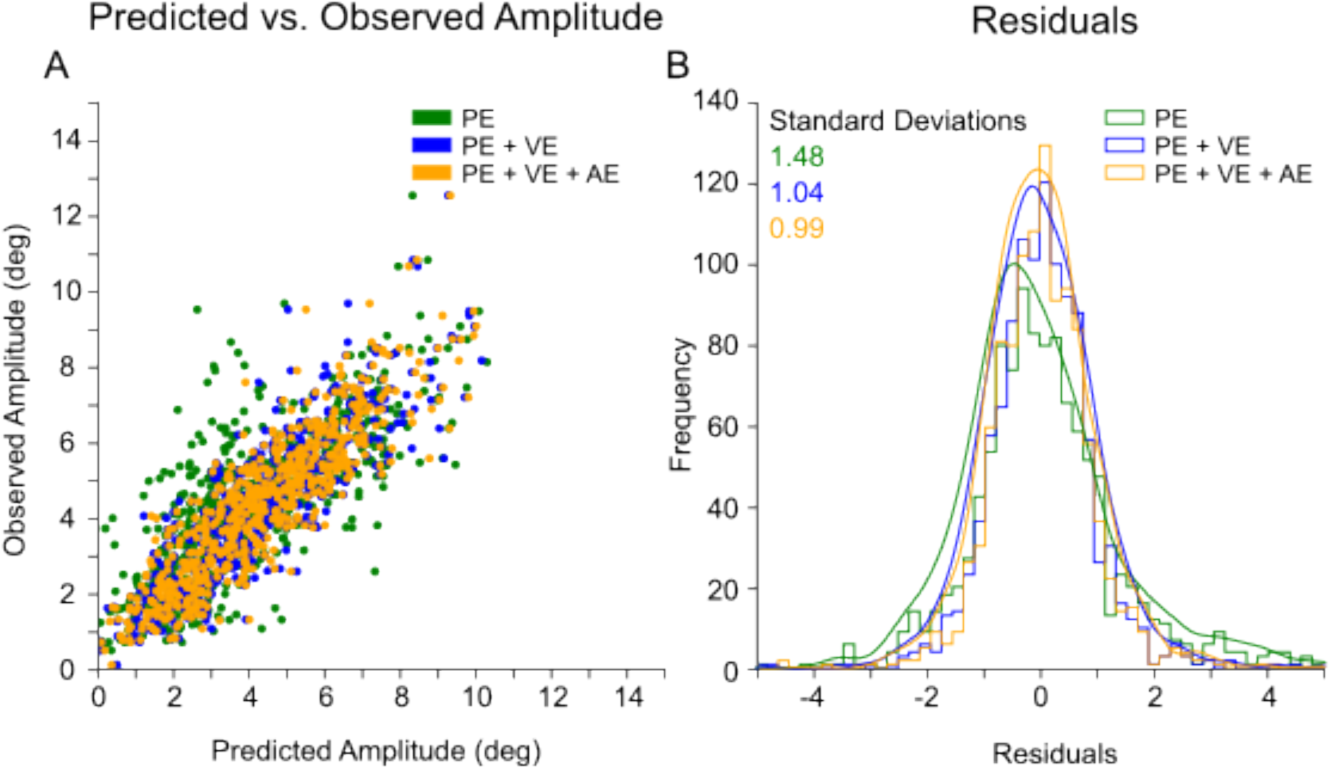
Visualization of how the regression model developed as each variable was added for an individual participant. A) Predicted vs. observed amplitude. Green: predicted vs. observed saccade amplitude values when position error was the only independent variable used for computing saccade amplitude. Blue: same with the addition of velocity error. Yellow: same including acceleration error. B) Histogram of residuals between predicted and observed saccade amplitudes, including kernel density estimates with each addition of a new variable. Residuals and standard deviations decrease with the addition of each variable. All analyses were performed using signals 100ms before saccade onset.

While retinal acceleration error was a significant predictor of saccade amplitude when all participants data was compiled and used in the analysis, there was some variability between individual participants (Table 2). When examined individually, retinal acceleration error was not a significant predictor of saccade amplitude in addition to position and velocity error in 5 of the 13 participants when computing saccade amplitude. The 8 other participants all reached statistical significance. Regression results for each individual participant are described below in table 2.

**Table 2.**
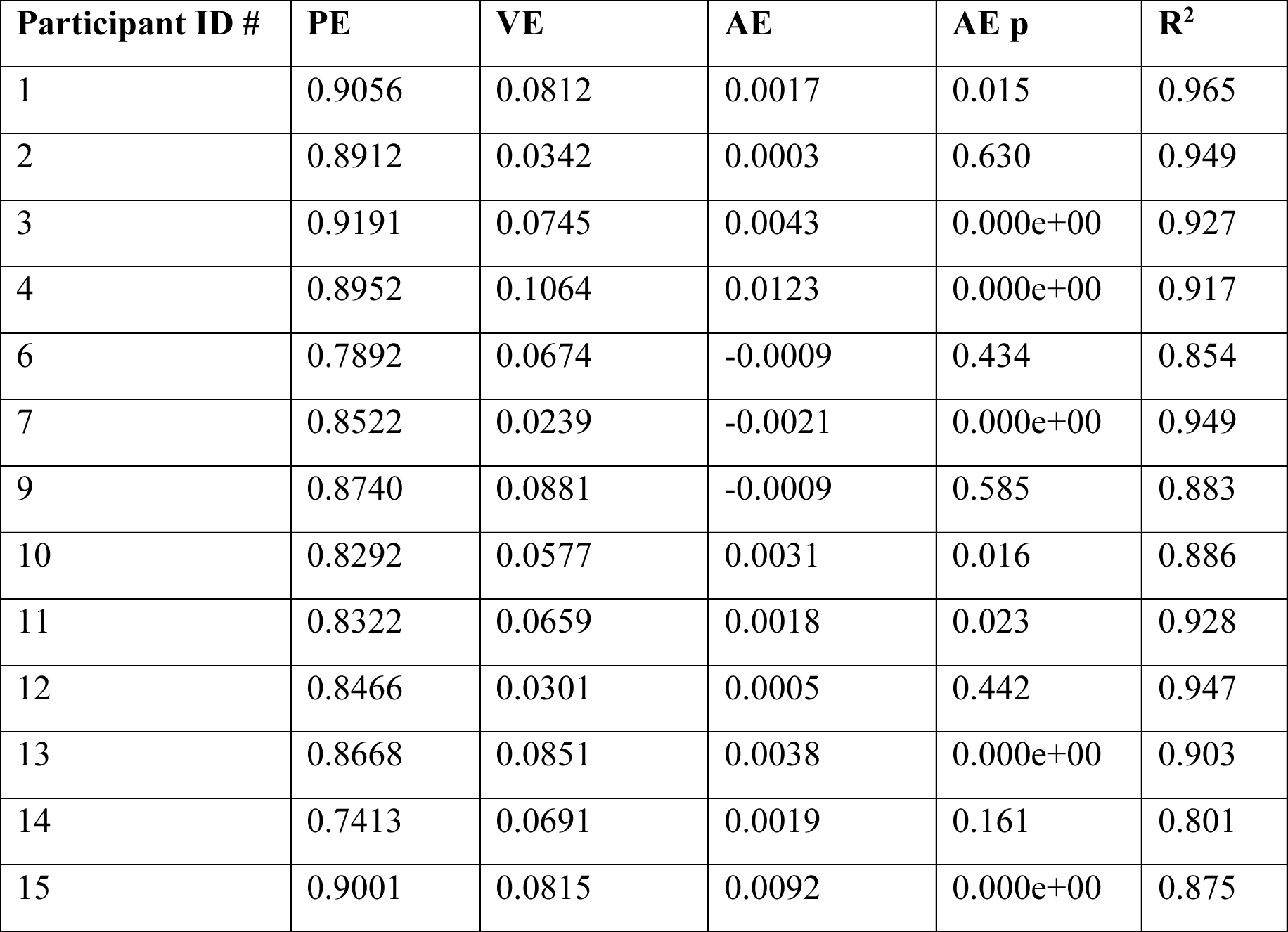
Multiple linear model regression results for individual participants.

In summary, our findings align with both our initial predictions and the existing body of literature on saccadic responses to accelerating target motion, as detailed in the Discussion section. Notably, our study highlights the relatively modest yet statistically significant contribution of retinal acceleration error in the computation of catch-up saccade amplitudes.

### Catch-up Saccade Timing

We also aimed to investigate the potential impact of retinal acceleration error on saccade timing. More specifically, our inquiry centered on whether retinal acceleration error would influence the predicted position error, and subsequently, saccade latencies. This would result in shorter saccade latencies when the predicted position error aligns with the direction of acceleration error and longer latencies when they oppose each other. This effect occurs because the addition or subtraction of the same acceleration magnitude to the predicted position error can enhance or diminish the certainty of the need for a saccade, therefore leading to shorter or longer saccade latencies depending on the relative sign.

Figure 4 illustrates the relationship between saccade latencies and predicted position error across different bin sizes, showing how they differ depending on the sign of acceleration error. In panels A and B, we can see that saccade latencies are shorter when predicted position errors are larger, as well as shorter when acceleration error has the same direction (sign) as predicted position error. This can be seen by a slight difference in the shape of latency distributions in panel A as well as by the difference in median latencies in panel B. Our interpretation is that these latencies are shorter due to there being less uncertainty as to whether a saccade should be triggered. Panel C illustrates the latency differences between the sign of acceleration error for each bin of predicted position error.

**Figure 4.**
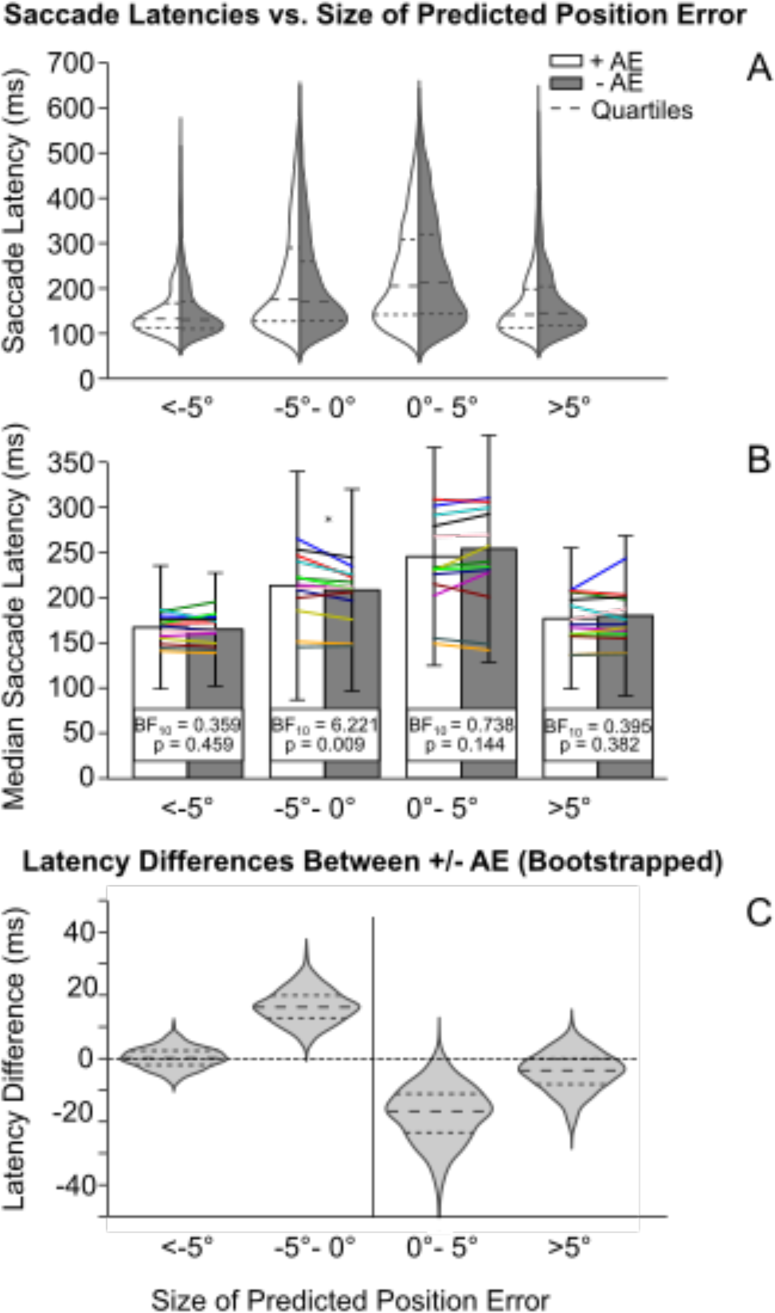
A) Saccade latency distributions of all participants across 4 bins of predicted position error split into groups of positive vs. negative retinal acceleration error. B) Median saccade latencies across all participants as well as for each individual participant (colored lines) . P-values and Bayes factor are from a paired t-test and a Bayesian paired t-test. Stars indicate statistical significance. C) Bootstrapped median saccade latency differences across all participants for each bin size of predicted position error.

We wanted to test whether latency differences for different sizes of predicted position error were statistically significant. To test this, we used a repeated-measures ANOVA with the sign of retinal acceleration error and predicted position error bin sizes as the independent variables and saccade latencies as the dependent variable. The ANOVA revealed a significant main effect of predicted position error (on saccade latencies F(3,36) = 40.975, p < .001), but no main effect of the sign of acceleration error (p>0.05). However, as expected there was a significant interaction between the effects of sign of acceleration error and size of predicted position error (F(3, 36) = 4.333, p = 0.010).

A paired sample t-test was performed to compare saccade latencies for each of the 4 predicted position error bins. There was a significant difference in saccade latencies between positive and negative retinal acceleration error or the −5 to 0 deg small negative predicted position error bin, (t(12)=3.101, p=0.009), but no significant difference for the three other bins < −5 deg large negative (t(12)=0.770, p=0.456), 0 to 5 deg small positive (t(12)=-1.562, p=0.144) and > 5 deg large positive (t(12)=-0.907, p=0.382).

A Bayesian paired sample t-test was also performed, revealing moderate evidence for a difference in saccade latencies between positive and negative acceleration error for small negative (−5 to 0 deg) predicted position errors (BF10 = 6.221), but no evidence for the other sizes of predicted position errors; < −5 deg large negative (BF10 = 0.359), 0 to 5 deg small positive (BF10 = 0.738) and > 5 deg large positive (BF10 = 0.395).

These results partially support our hypotheses; we did not expect the sign of acceleration error to have a main effect on saccade latencies since it averages out across all predicted position errors. However, we did expect a differential effect of sign of AE as a function of the predicted position error. If AE did not play a role in the timing of catch-up saccades, we would expect to see identical latency distributions for both positive and negative AEs. The results displayed in Figure 4 show otherwise. Predicted position error either increased or decreased in the same direction as the sign of AE. We can also observe that saccade latencies are shorter when the predicted position error is the same direction as AE for small predicted position error bins (though this did not reach significance for small positive predicted position errors). These findings affirm prior research on catch-up saccade triggering; longer saccade latencies were associated with smaller predicted position errors, while shorter latencies corresponded to larger predicted position errors (Nachmani et al., 2020).

Finally, we wanted to further analyze the effects of both retinal velocity and acceleration on saccade latencies across a finer scale of position errors at target step. Figure 5 displays saccade latencies vs position error at the target step binned both by size of velocity error (panel A) and acceleration error (panel B). In panel A, data is binned by size of velocity errors. Large negative is <-10 deg/s, small negative is −10 to 0 deg/s, small positive is 0 to 10 deg/s, and large positive is >10 deg/s. Panel B is binned by size of acceleration errors. Large negative is <-20 deg/s^2^, small negative is −20 to 0 deg/s^2^, small positive is 0 to 20 deg/s^2^, and large positive is >20 deg/s^2^. In both analyses, we see that saccade latencies are longest when position errors are smallest due to increased uncertainty as to whether a saccade is required. We also see similar results to figure 4 and statistical analyses – latencies are shorter when position errors are in the same direction of acceleration and velocity errors. This result is particularly clear for larger values of acceleration and velocity errors. The pattern of latencies shown in panel A corroborates the results of Nachmani et al. (2020), while panel B confirms the latter findings as well as our current study’s saccade timing hypothesis that saccade latencies are shorter when position error is in the same direction as acceleration error and vice versa. There is a less pronounced effect when binned by acceleration error rather than velocity error. This aligns with our expectations, considering the smaller relative impact of acceleration error, as outlined in the kinematic equation described in our timing hypothesis (Hyp. 2). This signed effect of retinal acceleration error on saccade latency is also correlated with saccade amplitude. Panel C displays a shift in latency distribution in that there are longer latencies when the sign of acceleration error is in the same direction as the sign of the amplitude.

**Figure 5.**
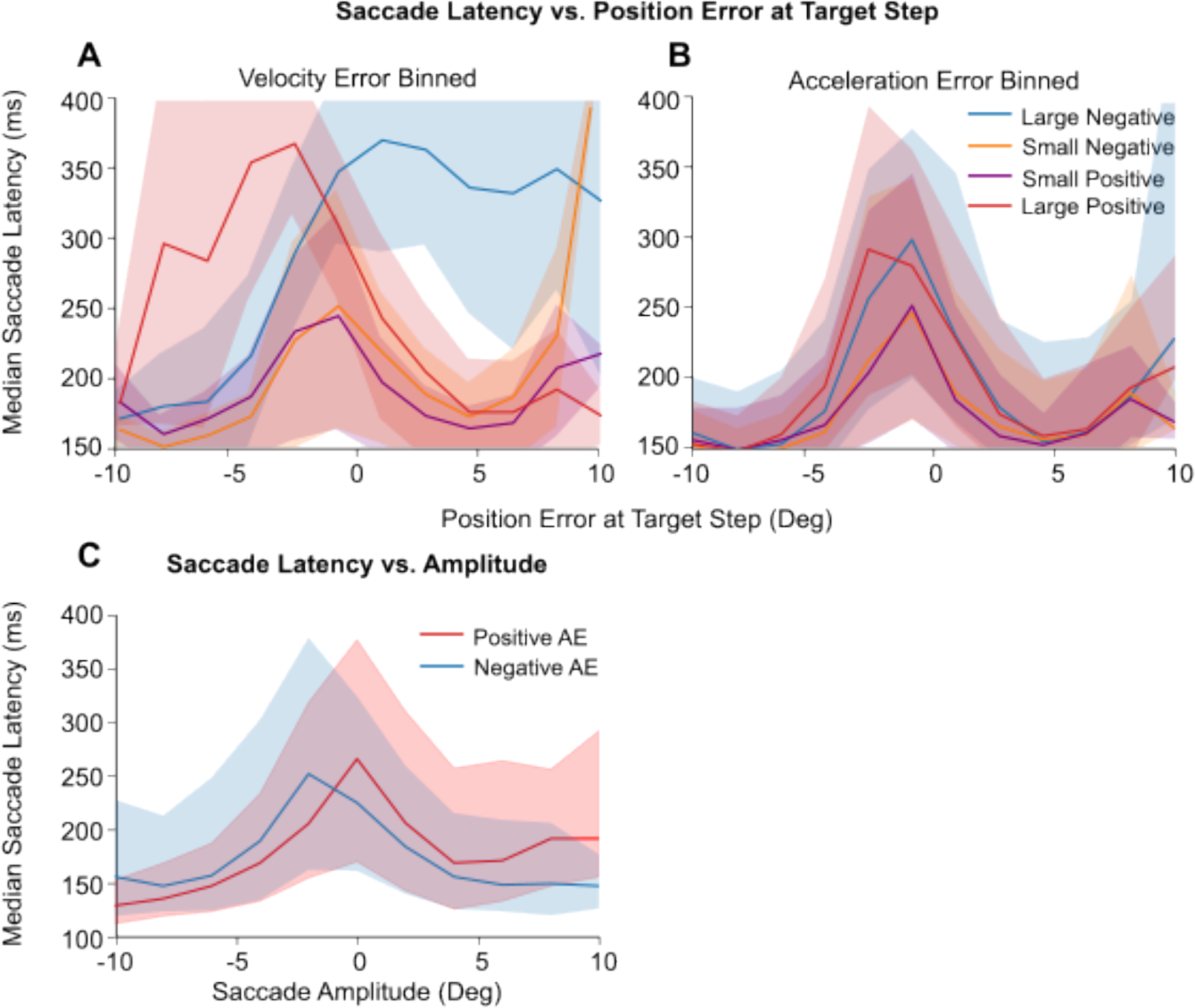
Median saccade latencies vs. position error and saccade amplitude. A) Saccade latencies as a function of position error at the step for different retinal velocity errors. Large negative:<-10 deg/s, small negative: −10 to 0 deg/s, small positive: 0 to 10 deg/s, large positive: >10 deg/s. B) Same as panel A but for different retinal acceleration errors. Large negative: <-20 deg/s^2^, small negative: −20 to 0 deg/s^2^, small positive: 0 to 20 deg/s^2^, large positive: >20 deg/s^2^. C) Median saccade latencies vs saccade amplitude binned by sign of acceleration error. Shaded areas display the 25^th^ – 75^th^ percentile range of the data.

## Discussion

### General Discussion

In this study, we investigated whether retinal acceleration error is used by the brain in determining the timing and amplitude of catch-up saccades during smooth pursuit. To test this, we designed an accelerating target tracking task and quantified the influence of retinal position, velocity and acceleration errors on the timing and amplitude of catch-up saccades during pursuit. We observed that retinal acceleration error influenced both the amplitude and timing of catch-up saccades, consistent with our hypotheses. In addition to position and velocity errors, acceleration error had a small influence on predicting saccade amplitude. Furthermore, when the direction of acceleration error and predicted position error was aligned, saccade latencies were shorter than when the signs opposed each other. These findings expand on the results of previous behavioural studies which demonstrated that humans are able to perceive and discriminate object acceleration (Bennett et al., 2007; Kreyenmeier et al., 2021; <u>Watamaniuk</u> et al., 2003). In summary, we provide evidence that the brain utilizes target acceleration information to compute the amplitude and timing of catch-up saccades.

### Comparison to the literature

The results of our study are confirmatory to others that investigate the computation of catch-up saccade amplitude (de Brouwer et al., 2002) and timing (Nachmani et al., 2020) in that we show that retinal position and velocity errors are used to predict catch-up saccade amplitude. Additionally, our study provides new evidence that retinal acceleration error is included in these computations. In comparison to the contribution of position and velocity errors, acceleration error has a small effect on predicting catch-up saccade amplitude. This can be explained by the scaling of our regression coefficients in eq 1. Indeed, given the temporal horizon of predicted position error, the expected amplitude correction that can be attributed to the position component is largest, followed by velocity, and acceleration being the smallest due to the nature of the kinematic equation.

In addition to this effect on catch-up saccade computations, acceleration has also been shown to influence the pursuit system. Kreyenmeier and colleagues’ (2022) recent occlusion study found that while acceleration is taken into account by the pursuit system leading up to target occlusion, it is not used in predicting time-to-contact when making an interceptive hand movement. These results suggest that acceleration signals are used differently when visually tracking compared to predicting manual interception.

The relatively small influence of acceleration error on saccade amplitude and timing could also be potentially linked to the difficulty humans have with perceiving acceleration, which has been a topic of debate. A prevalent theory posits that acceleration might be perceived indirectly through changes in speed (Bennett, de Xivry, Barnes & Lefèvre, 2007; Watamaniuk and Heinen, 2003). This conclusion is supported by findings that the participants can perceive accelerating target movement based on a threshold of 25% change in velocity throughout a trial (Brouwer, Brenner and Smeets, 2002).

### Hypothetical Neural Mechanisms

The oculomotor response to accelerating target motion has also been investigated from a neural perspective. Primate area MT is one of the core motion processing areas of the brain. Evidence from electrophysiology studies from MT neurons in monkeys shows joint target acceleration and target speed coding (Lisberger & Movshon 1999; Price et al. 2005). These studies support theories that suggest acceleration is processed by the brain indirectly through changes in velocity. Furthermore, frontal eye field (FEF) activity has been shown to reflect a dynamic internal representation of target motion (Barborica & Ferrera, 2003, 2004; Xiao, Barborica, & Ferrera, 2007). It is possible that retinal acceleration error could be processed by one or both of these motion processing areas in the brain, either directly or indirectly.

### Limitations

A limitation of the present study, shared with many eye-tracking investigations conducted in controlled laboratory settings, is the challenge of generalizing findings to real-world conditions. In our study, participants were tasked with following a sparse stimulus, a dot that moved unpredictably across a restricted range of accelerations, limited by screen dimensions. This stimulus lacked salience for the observers and its simplicity limited the development of prior expectations akin to those formed with natural stimuli. Research has shown that the human visual system responds differently when processing dynamic, natural scenes as opposed to simplified artificial stimuli typically used in laboratory experiments (Kayser, Körding & König, 2004). Indeed, the categorization of conventional laboratory stimuli demands more attentional resources compared to the relatively effortless processing of natural stimuli (Li, VanRullen, Koch & Perona, 2002). Furthermore, humans learn and develop priors related to the laws of motion throughout their lifetime when interacting with natural stimuli - for example with gravity (Zago and Lacquaniti, 2005; Jörges and López-Moliner, 2017). For example, in a naturalistic occlusion task portraying a virtual baseball game, humans track and predict fly-ball trajectories more accurately with natural gravity compared to manipulated gravity (Bosco et al., 2012). Together, these findings highlight the fact that the human visual system is adapted to the properties of its everyday input, and therefore can only be fully understood within a naturalistic context.

### Future Directions

Accounting for the influence of retinal acceleration error when designing target tracking tasks could be useful for various subfields of neuroscience. Our findings that catch-up saccades use retinal acceleration error independently of velocity and position errors can be studied more specifically from a modeling perspective. Specifically, models of catch-up saccades could now be updated to account for an independent influence of target acceleration involved in computing saccade amplitude and timing.

Next, we should investigate how retinal acceleration affects saccades to natural stimuli, allowing for more generalizable interpretations consistent with everyday life. Specifically, how does the brain take accelerating object motion in naturalistic environments into account, and how is this different than in laboratory environments? How does the oculomotor system differ in computing saccade timing and amplitude in natural scenes vs. simple, artificial stimuli. For example, would the target acceleration have more of an influence on saccade timing and amplitude when observing naturally falling objects, accelerating cars, or running humans and animals compared to a simple dot accelerating across a screen? Naturalistic research will be useful in updating current models of catch-up saccade behavior and better understanding their function in everyday life.

### Conclusion

Our study provides evidence that retinal acceleration error is used to compute the timing and amplitude of catch-up saccades to accelerating target motion in addition to retinal position and velocity errors. As expected, we found that the influence of retinal acceleration error on predicting catch-up saccade amplitude was small. We also found a signed effect of retinal acceleration error on saccade timing, in that latencies were shorter when retinal acceleration error and predicted position error were the same sign and longer when opposite.

## Acknowledgments

This work is supported by Natural Sciences and Engineering Research Council of Canada and Canada Foundation for Innovation. SD was supported by a Queen’s FHS Dean’s Graduate Award. Thank you to the members of the Blohm lab for your feedback.

## Notes

### Competing Interest Statement

The authors have declared no competing interest.

## References

Barborica, A., & Ferrera, V. P. (2003). Estimating invisible target speed from neuronal activity in monkey frontal eye field. Nature neuroscience, 6(1), 66–74.

Barborica, A., & Ferrera, V. P. (2004). Modification of saccades evoked by stimulation of frontal eye field during invisible target tracking. Journal of Neuroscience, 24(13), 3260–3267.

Bennett, S. J., & Barnes, G. R. (2006). Combined smooth and saccadic ocular pursuit during the transient occlusion of a moving visual object. Experimental brain research, 168, 313–321.

Bennett, S. J., & Benguigui, N. (2013). Is acceleration used for ocular pursuit and spatial estimation during prediction motion?. PLoS One, 8(5), e63382.

Bennett, S. J., de Xivry, J. J. O., Barnes, G. R., & Lefevre, P. (2007). Target acceleration can be extracted and represented within the predictive drive to ocular pursuit. Journal of Neurophysiology, 98(3), 1405–1414.

Bosco, G., Delle Monache, S., & Lacquaniti, F. (2012). Catching what we can’t see: manual interception of occluded fly-ball trajectories. PLoS One, 7(11), e49381.

Brainard DH (1997) The psychophysics toolbox. Spat Vis 10:433– 436.

Brostek, L., Eggert, T., & Glasauer, S. (2017). Gain control in predictive smooth pursuit eye movements: evidence for an acceleration-based predictive mechanism. eneuro, 4(3).

Coutinho, J. D., Lefèvre, P., & Blohm, G. (2021). Confidence in predicted position error explains saccadic decisions during pursuit. Journal of Neurophysiology, 125(3), 748–767.

De Brouwer, S., Missal, M., Barnes, G., & Lefèvre, P. (2002). Quantitative analysis of catch-up saccades during sustained pursuit. Journal of neurophysiology, 87(4), 1772–1780.

De Brouwer, S., Yuksel, D., Blohm, G., Missal, M., & Lefèvre, P. (2002). What triggers catch-up saccades during visual tracking?. Journal of neurophysiology, 87(3), 1646–1650.

Jörges, B., & López-Moliner, J. (2017). Gravity as a strong prior: implications for perception and action. Frontiers in Human Neuroscience, 11, 203.

Kayser, C., Körding, K. P., & König, P. (2004). Processing of complex stimuli and natural scenes in the visual cortex. Current opinion in neurobiology, 14(4), 468–473.

Keller, E., & Johnsen, S. S. (1990). Velocity prediction in corrective saccades during smooth-pursuit eye movements in monkey. Experimental Brain Research, 80, 525–531.

Krauzlis, R. J. (2004). Recasting the smooth pursuit eye movement system. Journal of neurophysiology, 91(2), 591–603.

Krauzlis, R. J., & Lisberger, S. G. (1989). A control systems model of smooth pursuit eye movements with realistic emergent properties. Neural Computation, 1(1), 116–122.

Krauzlis, R. J., & Lisberger, S. G. (1994). A model of visually-guided smooth pursuit eye movements based on behavioral observations. Journal of computational neuroscience, 1(4), 265–283.

Kreyenmeier, P., Kämmer, L., Fooken, J., & Spering, M. (2022). Humans can track but fail to predict accelerating objects. Eneuro, 9(5).

Li, F. F., VanRullen, R., Koch, C., & Perona, P. (2002). Rapid natural scene categorization in the near absence of attention. Proceedings of the National Academy of Sciences, 99(14), 9596–9601.

Lisberger, S. G., Evinger, C., Johanson, G. W., & Fuchs, A. F. (1981). Relationship between eye acceleration and retinal image velocity during foveal smooth pursuit in man and monkey. Journal of Neurophysiology, 46(2), 229–249.

Lisberger, S. G., Morris, E. J., & Tychsen, L. (1987). Visual motion processing and sensory-motor integration for smooth pursuit eye movements. Annual review of neuroscience, 10(1), 97–129.

Lisberger, S. G., & Movshon, J. A. (1999). Visual motion analysis for pursuit eye movements in area MT of macaque monkeys. Journal of Neuroscience, 19(6), 2224–2246.

Nachmani, O., Coutinho, J., Khan, A. Z., Lefèvre, P., & Blohm, G. (2020). Predicted position error triggers catch-up saccades during sustained smooth pursuit. eneuro, 7(1).

Orban de Xivry JJ, Lefèvre P (2007) Saccades and pursuit: two outcomes of a single sensorimotor process. J Physiol 584:11–23.

Price, N. S., Ono, S., Mustari, M. J., & Ibbotson, M. R. (2005). Comparing acceleration and speed tuning in macaque MT: physiology and modeling. Journal of Neurophysiology, 94(5), 3451–3464.

Rashbass, C. (1961). The relationship between saccadic and smooth tracking eye movements. The Journal of physiology, 159(2), 326.

Robinson, D. A. (1986). The systems approach to the oculomotor system. Vision research, 26(1), 91–99.

Tavassoli, A., & Ringach, D. L. (2010). When your eyes see more than you do. Current Biology, 20(3), R93–R94.

Xiao, Q., Barborica, A., & Ferrera, V. P. (2007). Modulation of visual responses in macaque frontal eye field during covert tracking of invisible targets. Cerebral Cortex, 17(4), 918–928.

Zago, M., & Lacquaniti, F. (2005). Visual perception and interception of falling objects: a review of evidence for an internal model of gravity. Journal of Neural Engineering, 2(3), S198.

